# Transcriptomic basis and evolution of ant nurse-larval social regulatory interactions

**DOI:** 10.1101/514356

**Authors:** Michael R. Warner, Alexander S. Mikheyev, Timothy A. Linksvayer

**Affiliations:** Department of Biology, University of Pennsylvania, Philadelphia, Pennsylvania, USA; Ecology and Evolution Unit, Okinawa Institute of Science and Technology, Onna, Okinawa, Japan; Research School of Biology, Australian National University, Canberra, ACT, Australia

## Abstract

Development is often strongly regulated by interactions among close relatives, but the underlying molecular mechanisms are largely unknown. In eusocial insects, interactions between caregiving worker nurses and larvae regulate larval development and resultant adult phenotypes. Here, we begin to characterize the social interactome regulating ant larval development by collecting and sequencing the transcriptomes of interacting nurses and larvae across time. We find that the majority of nurse and larval transcriptomes exhibit parallel expression dynamics across larval development. We leverage this widespread nurse-larva gene co-expression to infer putative social gene regulatory networks acting between nurses and larvae. Genes with the strongest inferred social effects tend to be peripheral elements of within-tissue regulatory networks and are often known to encode secreted proteins. This includes interesting candidates such as the nurse-expressed *giant-lens*, which may influence larval epidermal growth factor signaling, a pathway known to influence various aspects of insect development. Finally, we find that genes with the strongest signatures of social regulation tend to experience relaxed selective constraint and are evolutionarily young. Overall, our study provides a first glimpse into the molecular and evolutionary features of the social mechanisms that regulate all aspects of social life.

**Author Summary:** Social interactions are fundamental to all forms of life, from single-celled bacteria to complex plants and animals. Despite their obvious importance, little is known about the molecular causes and consequences of social interactions. In this paper, we study the molecular basis of nurse-larva social interactions that regulate larval development in the pharaoh ant *Monomorium pharaonis*. We infer the effects of social interactions on gene expression from samples of nurses and larvae collected in the act of interaction across a developmental time series. Gene expression appears to be closely tied to these interactions, such that we can identify genes expressed in nurses with putative regulatory effects on larval gene expression. Genes which we infer to have strong social regulatory effects tend to have weak regulatory effects within individuals, and highly social genes tend to experience relatively weaker natural selection in comparison to less social genes. This study represents a novel approach and foundation upon which future studies at the intersection of genetics, behavior, and evolution can build.

## Introduction

Social interactions play a prominent role in the lives of nearly all organisms [1] and strongly affect trait expression as well as fitness [2–4]. Social interactions in the context of development (e.g. parental care) often strongly regulate developmental trajectories and resultant adult phenotypes, for example via transferred compounds such as milk in mammals [5,6], milk-like secretions in arthropods [7,8], and other forms of nutritional provisioning [9,10]. In many taxa including certain birds, mammals, and insects, care for offspring and the regulation of offspring development has shifted at least in part from parents to adult siblings, who perform alloparental care [11]. In eusocial insect societies, sterile nurse workers regulate the development of their larval siblings by modulating the quantity and quality of nourishment larvae receive [12–14], as well as through the direct transfer of growth-regulating hormones and proteins [15,16]. At the same time, larvae influence nurse provisioning behavior via pheromones [17–20] and begging behavior [21,22].

In general, traits such as caregiving behavior that are defined or influenced by social interactions are the property of the genomes of multiple interacting social partners [2,14]. This has implications for both the mechanistic (e.g., molecular) underpinnings of development and trait expression as well as the genetic basis of trait variation at the population level -- i.e. how allelic variation in the genomes of interacting social partners affects trait variation [2,14]. Furthermore, because social traits are expressed in one individual but impact the fitness of other individuals, social behavior and socially-influenced traits experience distinct forms of selection, including kin selection and social selection [23,24]. Altogether, these distinct genetic features and patterns of selection are often thought to lead to distinct evolutionary features, such as rapid evolutionary dynamics in comparison to other traits [25–27]. In eusocial insects, previous studies show that variation in larval developmental trajectories and ultimate adult phenotypes (including reproductive caste, body size, etc.) depends on the combination of larval and nurse genotypes [28–34]. However, the identity of specific genes and molecular pathways that are functionally involved in the expression of social interactions (e.g., genes underlying nurse and larval traits affecting nurse-larva interactions) and the patterns of molecular evolution for these genes have remained less well studied [15,16,35,36].

Transcriptomic studies are often used to identify sets of genes underlying the expression of particular traits by performing RNA-sequencing on individuals that vary in the expression of such traits. For example, in social insects, recent studies have compared the transcriptomes of workers that perform nursing versus foraging tasks [37–39], or nurses feeding larvae of different stages or castes [35,40]. However, given the phenotypic co-regulation known to occur between interacting social partners (here, nurses and larvae), it is likely that genes expressed in one social partner affect the expression of genes in the other social partner, and vise-versa, such that interacting social partners are connected by “social” gene regulatory networks [14,32,41,42]. Thus, identifying the genes important for *social interactions* such as nurse-larva interactions is only possible by studying the transcriptomic dynamics of both interacting social partners across a time series of interactions.

To understand the transcriptomic basis of host-symbiont interactions, recent studies have reconstructed gene regulatory networks acting between hosts and symbionts by collecting and profiling the transcriptomes of each social partner across a time series of interactions [43–47]. Here, we use analogous methodology to study transcriptomic signatures of nurse-larva interactions in the pharaoh ant, *Monomorium pharaonis*. We sample a developmental time series of larvae as well as the nurses that feed each larval stage in this series, collecting individuals at the moment of interaction in order to identify genes involved in the expression of nurse-larva interactions, as well as genes affected by these interactions (i.e. the full “social interactome” [14]). Pharaoh ant nurses tend to specialize on feeding young versus old larvae, and nurses feeding young versus old larvae show different transcriptomic profiles [40]. Larval transcriptomic profiles also change over development [48,49]. Given these results, we predicted that we would observe concerted changes in broad-scale gene expression in larvae and their nurses across larval development (Fig 1), reflective of the functional importance of nurse-larva interactions. Based on our dual RNA-seq data, we infer social gene regulatory networks acting between nurses and larvae to identify candidate genes predicted to have important social regulatory effects. Finally, we combine our measures of social regulatory effects with available population genomic data [48] to characterize the patterns of molecular evolution of genes underlying nurse-larva interactions.

**Fig 1.**
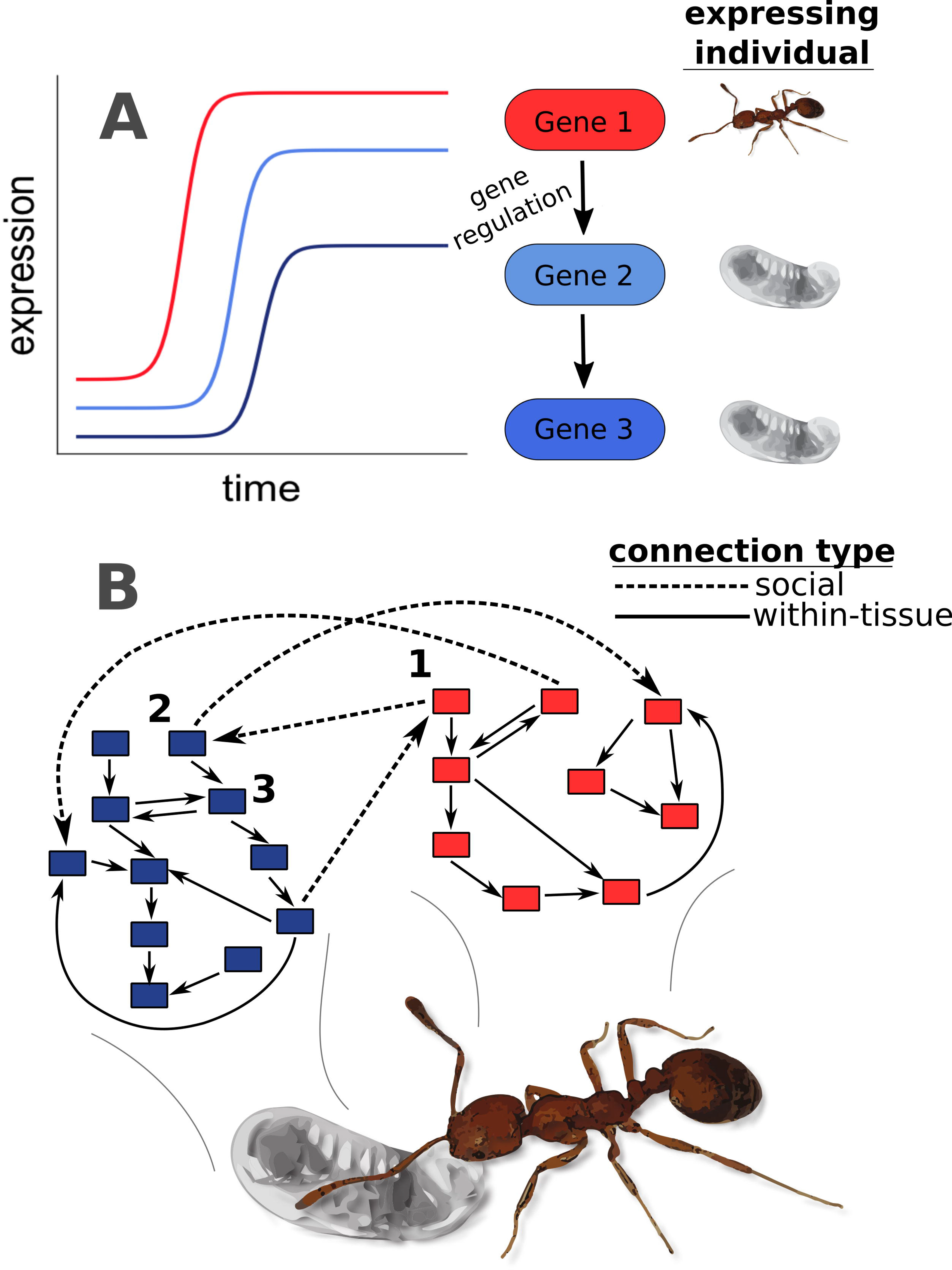
Social regulation of gene expression between ant nurses and larvae. (A) Cartoon depicting positive gene regulation (i.e. activation) between larvae and nurses, where gene 1 is expressed in nurses and genes 2 and 3 are expressed in larvae. After the expression of gene 1 increases, the expression of gene 2 increases as a result of the social interaction of nursing (depicted in [B]). This can occur if gene 1 itself codes for a protein passed to larvae, if the mRNA transcript is passed directly, or if gene 1 activates the expression of some other gene in nurses, which in turn is passed as mRNA (or codes for a protein that is passed) to larvae. Following the increase in expression of gene 2, the expression of gene 3, which is shown to be activated by gene 2, also increases. While we have depicted a time-lag in this social regulation of gene expression, the time lags are likely too short to observe in our data, as larvae were collected every 3-4 days across development. Therefore, correlated transcriptome dynamics over development (see Fig 2) would reflect mechanisms shown here. (B) Gene regulatory networks act between and within individuals engaged in social interactions. Blue boxes are genes expressed in larvae, and red boxes are genes expressed in nurses. Solid lines depict regulatory interactions within tissues (here, within larvae or within nurses), while dashed lines represent social connections (nurse-larva or vice versa).

## Results

### Transcriptome-wide signatures of nurse-larva co-expression across larval development

To elucidate transcriptomic signatures of nurse-larva interactions, we performed RNA-sequencing on worker-destined larvae across five developmental stages and nurses that fed larvae of each developmental stage (termed “stage-specific” nurses; see S1 Fig for sampling scheme, S1 Table for list of samples), building upon a previously published dataset focused on caste development in *M. pharaonis* [48]. We hypothesized that if genes expressed in larvae regulate the expression of genes in nurse and vice versa, we would observe correlated expression profiles across larval development in larvae and nurses (Fig 1). As a biological control, we collected “random nurses” that we observed feeding any stage of larvae in the colony, and hence would not be expected to show correlated expression dynamics with larvae across the five larval developmental stages. We also collected reproductive-destined larvae, but unless clearly stated otherwise, all analyses were performed on only worker-destined larvae. We collected ten individuals of each sample type to pool into one sample, and we sequenced whole bodies of larvae but separated nurse heads and abdomens prior to sequencing.

We grouped genes into co-expression profiles or “modules” using an algorithm designed to characterize gene co-expression dynamics across a short time series [50], known as Short Time-Series Expression Mining (STEM) [51]. Each module represents a standardized pre-defined expression profile, consisting of five values that each represent the log_2_ fold-change between the given developmental stage and the initial (L1) stage (see S2 Fig; this results in a total of 81 possible modules). We sorted genes into the module that most closely represented their expression profile by Pearson correlation. We identified modules containing a greater than expected number of genes, where we formed null expectations using permutation tests across developmental stages [50]. We identified such significantly-enriched modules separately for larvae, stage-specific nurse heads, stage-specific nurse abdomens, random nurse heads, and random nurse abdomens. We focused on both parallel (i.e. positive regulation or activation) and anti-parallel (i.e. inhibitory) correlated expression patterns by identifying significantly-enriched modules that were shared in both larvae and nurses (parallel), as well as significantly-enriched modules for which the inverse of the module was identified as significantly-enriched in the social partner (anti-parallel).

Larvae and stage-specific nurses shared many significantly-enriched modules (S2 Table). These shared modules contained the majority of genes expressed in nurses (65% of genes in stage-specific nurse heads and 76% in abdomens). A substantial proportion of the larval transcriptome was also shared with stage-specific nurse heads (22% of larval genes) and abdomens (60% of larval genes). Overall there was a widespread signature of correlated transcriptional patterns between stage-specific nurses and larvae across larval development (Fig 2A-D). These coordinated dynamics were dominated by parallel associations in nurse abdomens (possibly reflecting shared metabolic pathways) but anti-parallel associations in nurse heads (possibly reflecting the social regulation of larval growth). In contrast to stage-specific nurses, random nurses (our biological control) shared few significantly-enriched modules with larvae (S2 Table), and modules shared between random nurses and larvae contained significantly fewer genes than modules shared between stage-specific nurses and larvae (Fig 2E; Wilcoxon test, P < 0.001 for all comparisons). Specifically, 2% of genes expressed in random nurse heads and 13% of genes expressed in random nurse abdomens were in modules shared with larvae; 3% of genes expressed in larvae were in modules shared with random nurse heads, and 2% of genes expressed in larvae were in modules shared with random nurse abdomens.

**Fig 2.**
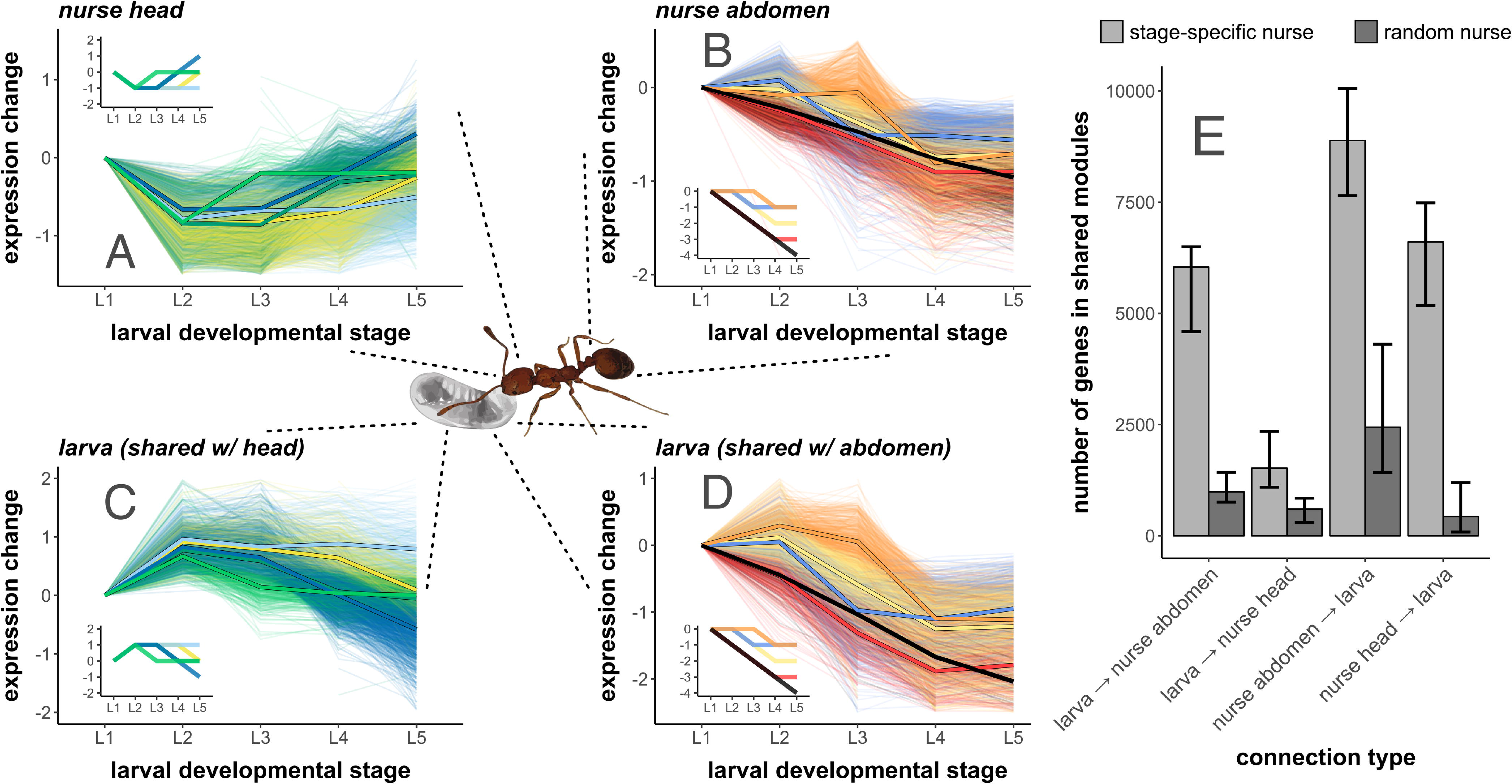
Nurse and larval transcriptomes show strong signatures of gene co-expression across larval development. Plots (A-D) depict the expression profiles of individual genes (light lines) as expressed in (A) nurse head, and (B) nurse abdomens, as well as (C) larvae, shared with nurse heads, and (D) larvae, shared with nurse abdomens. Dark lines indicate the median expression values of all genes sorted into modules, with pre-defined expression profiles of modules depicted in plot insets. Colors indicate the pre-defined expression profile (i.e. module) that genes have been sorted into. Only the five shared modules containing the most nurse-expressed genes are shown for clarity. Larval expression profiles are divided by the nurse tissue they are shared with, such that (C) depicts larval gene expression shared with nurse heads (A), while (D) depicts larval gene expression shared with nurse abdomens (B). Note that nurse heads and larvae shared inversely-related expression profiles, and that this algorithm does not reveal the direction of regulation as it is simply correlation-based. (E) Stage-specific nurses have more genes than random nurses in modules shared with larvae than do random nurses, reflecting more broad-scale co-expression across development. “Connection type” refers to the tissue that the number of genes was calculated in (i.e. larva → nurse head indicates the number of genes expressed in larvae that are in modules shared with nurse heads), though directionality is not determined in this algorithm. Error bars indicate 95% confidence intervals derived from systematic drop-1 jackknifing of nurse samples. N = 10944 genes total.

### Identification of genes putatively involved in social interactions

Given that we observed transcriptome-wide patterns consistent with nurse-larva transcriptional co-regulation across larval development, we next identified the genes that might be driving these patterns (see S3 Fig). We performed differential expression analysis to identify genes that varied in larval expression according to larval developmental stage, as well as genes that varied in nurse expression according to the developmental stage of larvae they fed. We identified 8125 differentially expressed genes (DEGs) in larvae (78% of 10446 total genes). We identified 2057 and 1408 DEGs in stage-specific nurse heads and abdomens, respectively, compared to 599 and 520 DEGs in random nurse heads and abdomens, respectively. We removed genes differentially expressed in both stage-specific and random nurses (N = 272 DEGs in heads, N = 140 DEGs in abdomens), which might differ among our colony replicates due to random colony-specific effects that were not consistently associated with social regulation of larval development. After this removal, we retained the top 1000 DEGs, sorted by P-value, for each sample type other than random nurses (larvae, stage-specific nurse heads, stage-specific nurse abdomens) for social gene regulatory network reconstruction, reasoning that these genes were the most likely to be involved in the regulation of larval development.

### Reconstruction of social gene regulatory networks

To infer putative gene-by-gene social regulatory relationships between nurses and larvae, we reconstructed gene regulatory networks acting within and between nurses and larvae (S3 Fig). The output of regulatory network reconstruction is a matrix of connection strengths, which indicate the regulatory effect (positive or negative) one gene has on another, separated according to the tissue the gene is expressed in. To identify the most highly connected (i.e. centrally located, upstream) genes of regulatory networks, we calculated within-tissue connectivity and social connectivity by averaging the strength of connections across each connection a gene made, differentiating between within-tissue (nurse-nurse or larva-larva) and social connections (nurse-larva) (Fig 1B). On average, within-tissue connectivity was higher than social connectivity (Wilcoxon test; P < 0.001 in all tissues), and within-tissue connectivity was negatively correlated with social connectivity in each tissue (S4 Fig). The top enriched gene ontology terms based on social connectivity in nurses were entirely dominated by metabolism (S3,S4 Tables; see also S5 Table for the top 20 genes by nurse social connectivity).

### Secreted proteins and social gene regulation

While based on our data it is not possible to distinguish between genes that code for protein products that are actually exchanged between nurses and larvae versus genes that affect behavior or physiology within organisms (Fig 1A), proteins that are known to be cellularly secreted represent promising candidates for the social regulation of larval development [40]. We downloaded the list of proteins that are known to be cellularly secreted from FlyBase [52] and used a previously-generated orthology map to identify ant orthologs of secreted proteins [40]. Genes coding for proteins with orthologs that are cellularly secreted in *Drosophila melanogaster* had higher social connectivity than genes coding for non-secreted orthologs in nurse heads (Fig 3A; Wilcoxon test; P = 0.025), though not for nurse abdomens (P = 0.067).

**Fig 3.**
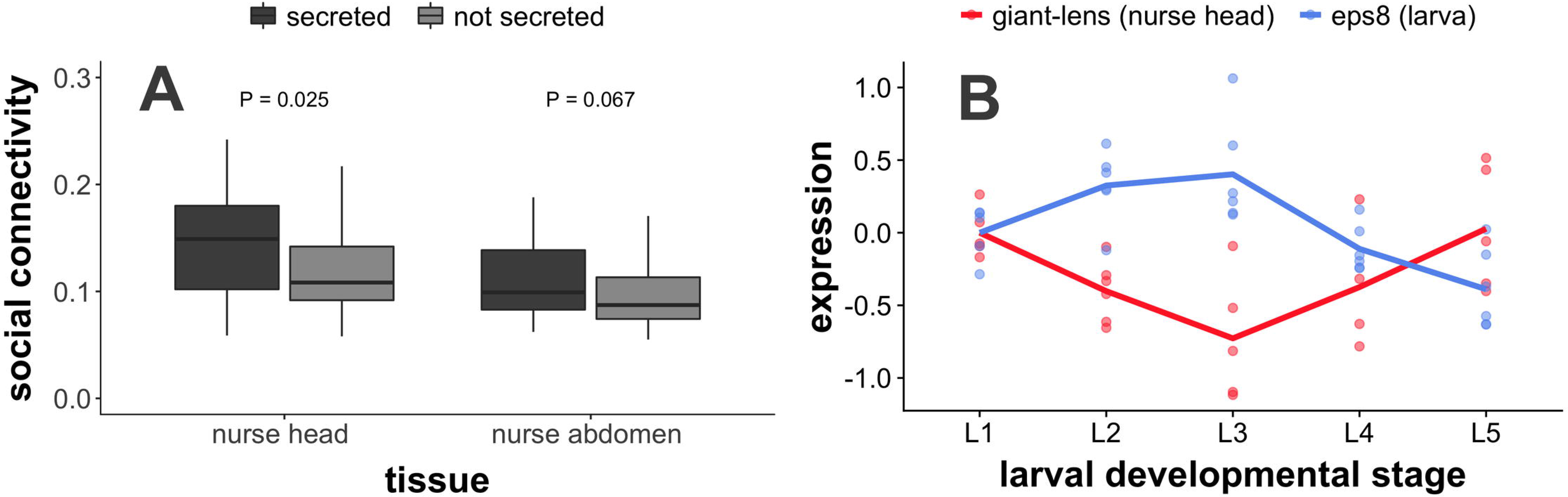
Genes encoding secreted proteins such as *giant-lens* are important for social gene regulation. (A) Genes encoding for proteins that are secreted in *Drosophila melanogaster* exhibit higher social connectivity (i.e. more strongly socially regulate larval expression) in nurse heads than genes encoding for non-secreted proteins (P-values from Wilcoxon test). (B) The protein *giant-lens* is one of the genes coding for secreted proteins with the highest social connectivity in nurse heads. Based on our data, *giant-lens* expressed in stage-specific nurse heads (red) appears to inhibit the expression of the homolog of human EGFR substrate 8 (*eps8*) expressed in worker-destined larvae (blue). The expression of *giant-lens* in nurses of a given colony was negatively correlated to the expression of *eps8* in larvae of the same sampled colony (rho = −0.270, P < 0.001, N = 25 colony/stage pairings after removing missing samples). Expression at stage *i* is equal to log_2_(expressioni/expression1), i.e. the ratio of expression at the given stage to expression at L1.

For the most part, we have focused on broad patterns of nurse-larva gene coregulation. In this paragraph, we will highlight the potential social role of one of the genes with the highest social connectivity within nurse heads, *giant-lens* (S6 Table; *giant-lens* is the 7^th^ highest gene coding for secreted proteins by social connectivity in nurse heads). *Giant-lens* is an inhibitor of epidermal growth factor receptor (EGFR) signaling [53], and g*iant-lens* expression in nurse heads was negatively associated with the expression of the homolog of *eps8*, human EGFR substrate 8 in larvae, most prominently seen in the spike in nurse *giant-lens* expression accompanied by a drop in larval *eps8* expression at the end of larval development (Fig 3B). *Giant-lens* was also used in regulatory network reconstruction in larvae (i.e. it was one of the top 1000 DEGs), and *giant-lens* expression in larvae drops steadily throughout development (S5 Fig; in contrast to the pattern of *giant-lens* expression in nurse heads). Interestingly, *eps8* does not exhibit a similar peak and drop in expression level in reproductive-destined larvae in comparison to worker-destined larvae (S6 Fig). It is important to note that these patterns were not seen for all genes in the EGFR pathway, and the results presented here cannot be taken as concrete evidence of EGFR regulation via social processes. Nonetheless, the mechanism illustrated here represents a tangible example of how nurse-larva interactions could function at the molecular level.

### Molecular evolution of social gene regulatory networks

To investigate the selective pressures shaping social regulatory networks, we used population genomic data from 22 resequenced *M. pharaonis* workers, using one sequenced *M. chinense* worker as an outgroup [48]. Using polymorphism and divergence data, we estimated gene-specific values of selective constraint, which represents the intensity of purifying selection that genes experience [54]. To identify genes disproportionately recruited to the core of social regulatory networks, we calculated “sociality index” as the difference between social connectivity and within-tissue connectivity for each gene. Sociality index was negatively correlated to selective constraint due to a positive correlation between within-tissue connectivity and constraint and a negative correlation between social connectivity and constraint (Fig 4A-C). Additionally, genes differed in sociality index according to their estimated evolutionary age, with ancient genes exhibiting lower sociality indices than genes in younger age categories (Fig 4D). Finally, while evolutionary age and evolutionary rate appear to be somewhat empirically confounded [55], selective constraint and evolutionary age were each independently associated with sociality index, based on a model including both variables as well as tissue (GLM; LRT; evolutionary age: χ^2^ = 21.536, P < 0.001; selective constraint: χ^2^ = 22.191, P < 0.001).

**Fig 4.**
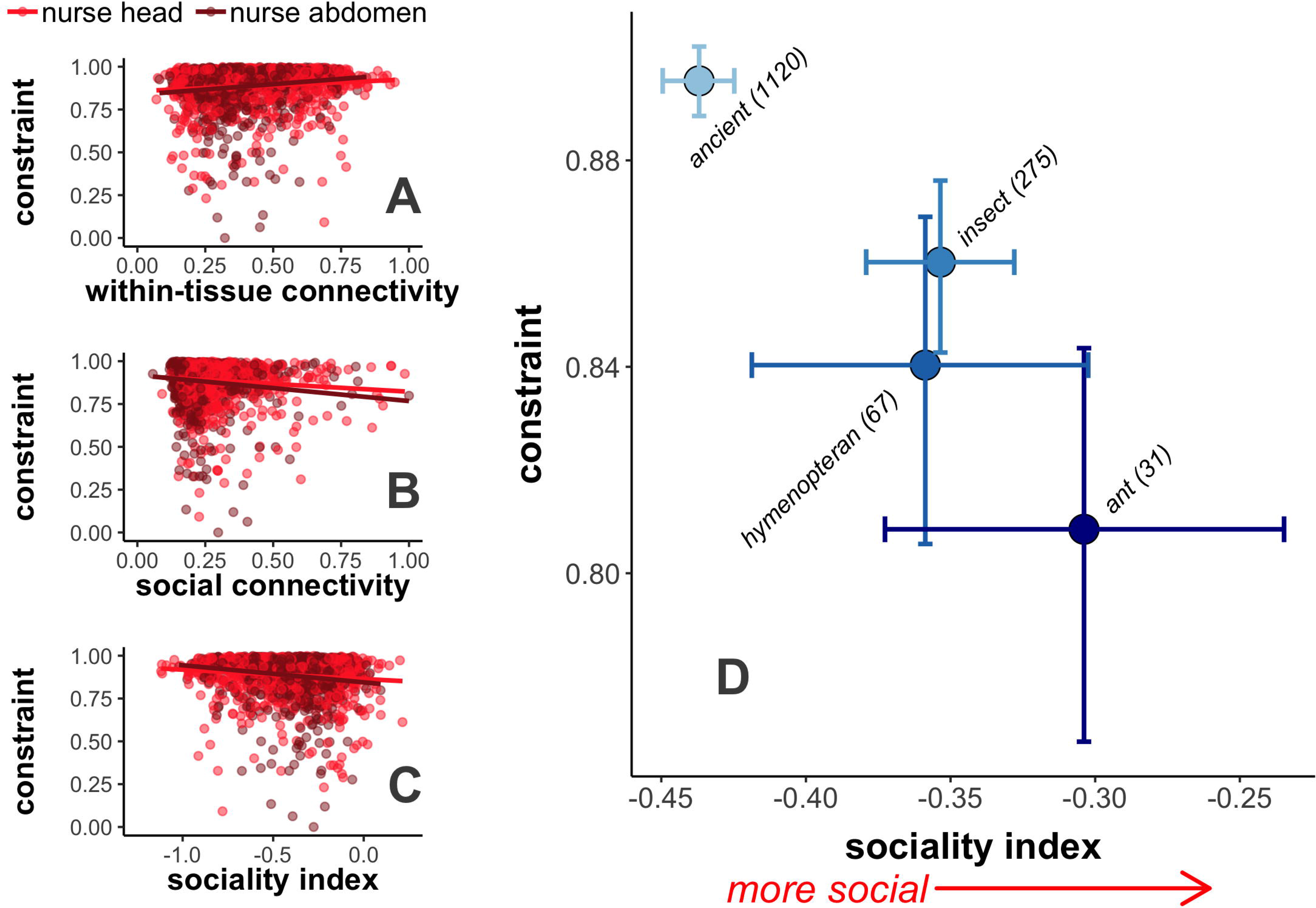
Highly social genes tend to be less evolutionarily constrained. Selective constraint, estimated from whole-genome polymorphism data, is (A) positively correlated with within-tissue connectivity (Spearman correlation; head: rho = 0.122, P < 0.001; abdomen: rho = 0.217, P < 0.001), but negatively correlated with (B) social connectivity (head: rho = −0.090, P = 0.009; abdomen: rho = −0.150, P < 0.001) and (C) sociality index (head: rho = −0.132, P < 0.001; abdomen: rho = −0.223, P < 0.001), where sociality index is the difference between social and within-tissue connectivity per gene. Each point in (A-C) indicates a single gene, as expressed in nurse heads or abdomens. Lines are trendlines from linear model. (D) Sociality index differs according to estimated evolutionary age (GLM; LRT; *χ*^2^ = 57.357, P < 0.001), as ancient genes tended to have lower sociality indices than all other categories (Tukey’s post-hoc test; ancient - insect: P < 0.001, ancient - hymenoptera: P < 0.001, ancient - ant: P < 0.001, all other comparisons P > 0.05). Individual points depict average values across nurse heads and abdomens for all genes within each estimated evolutionary age class, indicated by labels on points. Error bars depict 95% confidence intervals from bootstrapping. Numbers in parentheses indicate number of genes in each age class.

## Discussion

In organisms with extended offspring care, developmental programs are controlled in part by socially-acting gene regulatory networks that operate between caregivers and developing offspring [14,42]. In this study, we sequenced the transcriptomes of ant nurses and larvae as they interacted across larval development to assess the effects of social interactions on gene expression dynamics. We found that large sets of genes (i.e. modules) expressed in ant larvae and their caregiving adult nurses show correlated changes in expression across development (Fig 2). The majority of nurse and larval transcriptomes was represented in these correlated modules, suggesting that the tight phenotypic co-regulation characterizing nurse-larva interactions over the course of larval development is also reflected at the molecular level.

To characterize the overall network and evolutionary patterns of genes involved in nurse-larva interactions, we reverse engineered nurse-larva gene regulatory networks and calculated the “social connectivity” for each gene, defined as the sum of inferred social regulatory effects on all genes expressed in social partners. We found that genes with high social connectivity tended to have low within-individual connectivity (S4 Fig; where within-individual connectivity is defined as the sum of inferred regulatory effects acting within a given tissue). Nurse-expressed genes with higher sociality indices (i.e disproportionately higher social connectivity than within-individual connectivity) tended to be evolutionarily young and rapidly evolving due to relaxed selective constraint (Fig 4). Genes with high social connectivity were enriched for a number of Gene Ontology (GO) categories associated with metabolism (S3,S4 Tables), consistent with the idea that molecular pathways associated with metabolism are involved in the expression of social behavior [56,57]. Previously, many of the proteins found to be widely present in social insect trophallactic fluid transferred from nurses to larvae were involved in sugar metabolism (e.g. Glucose Dehydrogenase, several types of sugar processing proteins) [15]. Along the same lines, many of the genes with with high social connectivity in our study are also annotated with terms associated with sugar metabolism (S5 Table; e.g. Glycerol-3-phosphate dehydrogenase, Glucose dehydrogenase FAD quinone, Pyruvate dehydrogenase). Finally, we found that genes encoding for orthologs of cellularly-secreted proteins in *Drosophila melanogaster* (possibly important for intercellular signaling) tended to exhibit higher levels of social connectivity than their non-secreted counterparts (Fig 3A).

One gene that stands out in terms of being cellularly secreted and exhibiting a relatively high social connectivity is *giant-lens*, which inhibits EGFR signaling [53]. EGFR signaling affects eye and wing development [58] as well as body size in *D. melanogaster* [59], caste development in the honey bee *Apis mellifera* [59,60] via the transfer of royalactin from nurses to larvae [59], and worker body size variation in the ant *Camponotus floridanus* [61]. Further experimental work is necessary to ascertain whether *giant-lens* is actually orally secreted by nurses and transferred to larvae, but gene expression dynamics are consistent with the social transfer of *giant-lens* from nurses to larvae, followed by the inhibition of EGFR signaling at the end of larval development in worker-destined larvae (Fig 3B). Importantly, this inhibition is not seen in reproductive-destined larvae (S6 Fig). While caste in *M. pharaonis* is socially regulated in the first larval stage [49], social inhibition of EGFR signaling could play a role in the regulation of worker body size [61] or secondary caste phenotypes such as wings [62,63].

In terms of broad evolutionary patterns, our study complements previous results suggesting genes with worker-biased expression tend to be rapidly evolving, evolutionarily young, and loosely connected in regulatory networks in comparison to genes with queen-biased expression [38,48,64–66]. Because pharaoh ant workers are obligately sterile, their traits are shaped indirectly by kin selection, based on how they affect the reproductive success of fertile relatives (i.e. queens and males) [23,67]. As a result, all-else-equal, genes associated with worker traits are expected to evolve under relaxed selection relative to genes associated with queen traits [68,69].

In general, the suite of genic characteristics commonly associated with worker-biased genes (rapidly evolving, evolutionarily young, loosely connected) are all consistent with relaxed selection acting on genes associated with workers [49]. Here, we show that within the worker caste, genes that appear to be functionally involved in the expression of social behavior (i.e. nursing) experience relaxed selective constraint relative to genes important for within-worker processes. Therefore, the combination of kin selection as well rapid evolution thought to be characteristic of social traits [25] likely act in concert to shape the labile evolutionary patterns commonly associated with worker-biased genes. Finally, it has also been suggested that plastic phenotypes such as caste recruit genes which were evolving under relaxed selection prior to the evolution of such plastic phenotypes [70–72]. Our results could also be consistent with this hypothesis, though the population genomic patterns we observe show that relaxed selective constraint is ongoing.

In this study, we sought to reconstruct regulatory networks acting between nurses and larvae, beginning with the assumption that nurse gene expression changes as a function of the larval stage fed. This is more likely to be the case when nurses are specialized on feeding particular larval stages. According to a previous study, about 50% of feeding events are performed by specialists (though note specialization is likely a continuous trait, and the 50% figure is the result of a binomial test) [40]. Therefore, we expect our stage-specific nurse samples to comprise about 50% specialists. We also expect random nurse samples to contain 50% specialist nurses, but, crucially, the specialists should be relatively evenly divided among larval stages since random nurses were collected regardless of which larval stage they were observed feeding. Because our stage-specific nurse samples did not consist of 100% specialists, we expect that the signal of nurse-larva co-expression in our analysis is effectively diluted. In order to maximize the signal of nurse-larval co-expression dynamics, future studies would ideally focus entirely on specialists, as well as on tissues such as brains and the specific exocrine glands [73] known to be important for social behavior and communication. Despite these limitations, we were still able to observe transcriptomic signatures consistent with the social regulation of larval development.

## Conclusions

In this study, we uncovered putative transcriptomic signatures of social regulation and identified distinct evolutionary features of genes that underlie “social physiology”, the communication between individuals that regulates division of labor within social insect colonies [74,75]. Because we simultaneously collected nurses and larvae over a time series of interactions, we were able to elucidate the putative molecular underpinnings of nurse-larval social interactions. This is a promising approach that could be readily extended to study the molecular underpinnings of all forms of social regulation in social insect colonies, including regulation of foraging, regulation of reproduction, etc.. Furthermore, by adapting the methodology presented here (i.e. simultaneous collection over the course of interactions followed by sequencing), the molecular mechanisms and evolutionary features of genes underlying a diverse array of social interactions, including courtship behavior, dominance hierarchy formation, and regulation of biofilm production could all be investigated. Overall, this study provides a foundation upon which future research can build to elucidate the genetic underpinnings and evolution of interacting phenotypes.

## Methods

This study builds on previous work investigating genomic signatures of kin selection in which we characterized transcriptomic profiles from adult queens and workers, as well as queen- and worker-destined larvae [48]. While stage-specific nurses were used in the previous analysis, the knowledge of the developmental stage of larvae they fed was not, as they were simply treated as adult workers. This study also complements the past dataset with new data from random nurses, which were collected concurrently with previous samples.

### Study Design

To construct experimental colonies, we began by creating a homogenous mixture of approximately fifteen large source colonies of the ant *Monomorium pharaonis*. From this mixture, we created thirty total replicate experimental colonies of approximately equal sizes (~300-400 workers, ~300-400 larvae). We removed queens from ½ the study colonies to promote the production of reproductive-destined larvae. Reproductive caste is determined in *M. pharaonis* by the end of the first larval instar, likely in the egg stage [76], and queen presence promotes culling of reproductive-destined L1 larvae. Removing queens halts this culling, but it is unknown which colony members actually perform such culling [76]. While we initially expected the presence of queens to impact the gene expression profiles of nurses, we detected 0 DEGs (FDR < 0.1) between queen-present and queen-absent colonies for every sample type. This could indicate that nurses don’t perform culling and that worker developmental trajectories (and nutritional needs) are not appreciably different between queen-present and queen-absent colonies. Because queen presence did not substantially impact gene expression, in this study we pooled samples across queen-present and queen-absent colonies for all analyses.

We pre-assigned colonies to one of five larval developmental stages (labeled L1-L5, where L1 and L2 refer to 1st-instar and 2nd-instar larvae and L3, L4, and L5 refer to small, medium, and large 3rd-instar larvae [77]). We identified larval stage through a combination of hair morphology and body size. L1 larvae are nearly hairless, L2 larvae have straight hairs and are twice the length of L1 larvae, and L3-L5 larvae have dense, branched hairs [78]. We separated 3rd-instar larvae into three separate stages based on body size [77] because the vast majority of larval growth occurs during these stages. We sampled individuals (larvae as well as nurses) across larval development time: beginning at the L1 stage, we sampled colonies assigned to each subsequent stage at intervals of 3-4 days, by the time the youngest larvae in colonies lacking queens were of the assigned developmental stage (note that in colonies lacking queens, no new eggs are laid so the age class of the youngest individuals progressively ages). We sampled each colony once, according to the developmental stage we had previously assigned the colony (e.g. for colonies that we labeled ‘L4’, we waited until it was time to sample L4 larvae and nurses and sampled individuals from that colony at that time). From each colony, we sampled stage-specific nurses and worker-destined larvae, as well as random nurses from colonies with queens and reproductive-destined larvae from colonies without queens (starting at the L2 stage, because at L1 caste cannot be distinguished [76,77]. Reproductive-destined larvae include both males and queens (which cannot be readily distinguished), though samples are expected to be largely made up of queen-destined individuals given the typically skewed sex ratio of *M. pharaonis* [48]. See S1 Table for full sample list.

For each time point in each assigned colony, we collected stage-specific nurses, nurses feeding larvae of the specified developmental stage (L1, L2, etc). Concurrently, we collected random nurses, nurses we observed feeding a larva of any developmental stage. Rather than paint-marking nurses, we collected them with forceps as soon as we saw them feeding larvae. We collected random nurses as soon as we observed them feeding a larva of any developmental stage in the course of visually scanning the colony. We did not make an attempt to systematically collect nurses from different areas of the nest but did so haphazardly, such that the distribution of larval stages fed resembled overall colony demography. Nurses feed L1 and L2 larvae exclusively via trophallaxis (i.e. liquid exchange of fluid), while nurses feed L3-L5 larvae both via trophallaxis and by placing solid food in larval mouthparts [79]. To get a representative sample of all types of nurses, we did not distinguish between nurses feeding liquid and solid food, though all L3-L5 samples contained a mixture of the two. After collecting nurses, we anaesthetized the colony using carbon dioxide and collected larvae of the specified developmental stage. All samples were flash-frozen in liquid nitrogen immediately upon sample collection. Note that workers in *M. pharaonis* are monomorphic [80].

We performed mRNA-sequencing on all samples concurrently using Illumina HiSeq 2000 at Okinawa Institute of Science and Technology Sequencing Center. Reads were mapped to the NCBI version 2.0 *M. pharaonis* assembly [38], and we used RSEM [81] to estimate counts per locus and fragments per kilobase mapped (FPKM) for each locus. For further details on RNA extraction and library preparation, see [48].

### Transcriptome-wide signatures of nurse-larva co-expression across larval development

We used an algorithm that categorizes genes based on their expression dynamics over time into a number of modules represented by pre-defined expression profiles [50]; see S2 Fig for workflow). To create modules, we started at 0 and either doubled, halved, or kept the expression level the same at each subsequent stage, resulting in 81 possible modules (3*3*3*3 = 81; four stages after L1). To generate gene-specific expression profiles based on real results, we calculated the average log_2_ fold change in expression (FPKM) of the gene at each developmental stage compared to the initial expression level at stage L1. We then assigned each gene to the closest module by Pearson correlation between gene expression profile and module expression profile [50]. To identify significantly-enriched modules, we generated null distributions of the number of genes present in each module (based on permutation of expression over time), and retained modules with a significantly greater than expected number of genes based on these null distributions (FDR < 0.05 after Bonferroni multiple correction [50]).

### Identification of genes putatively involved in social interactions

We used the package EdgeR [82] to construct models including larval developmental stage and replicate and performed differential expression analysis for each sample type separately. We retained genes differentially expressed according to a nominal P-value of less than 0.05 (i.e. no false discovery correction), as the purpose of this step was simply to identify genes that could be involved in interactions that shape larval development (rather than spurious interactions arising from replicate-specific effects). See S1 Dataset for a list of all stage-specific nurse and larval differentially expressed genes.

### Social regulatory network reconstruction

We normalized expression for each gene using the inverse hyperbolic sine transformation of FPKM. As input to the algorithm, we constructed “meta-samples” by combining expression data within the same replicate and time point from nurses and larvae and labeling genes according to the tissue they were expressed in, along the lines of host-symbiont studies [43,45]. We utilized the program GENIE3 [83,84] to construct two types of networks: those acting between larvae and nurse heads, and those acting between larvae and nurse abdomens.

GENIE3 uses a random forest method to reconstruct regulatory connections between genes, in which a separate random forest model is constructed to predict the expression of each gene, with the expression of all other genes as predictor variables. The output of GENIE3 is a matrix of pairwise directional regulatory effects, where the regulatory effect of gene *i* on gene *j* is estimated as the feature importance of the expression of gene *i* for the random forest model predicting the expression of gene *j* (i.e. regulatory effect is how important the expression of gene *i* is for determining the expression of gene *j*). These regulatory effects (or strengths) include both positive and negative as well as non-linear effects, though these different effect types are not distinguished.

As a side note, a version of GENIE3 that was developed for time series data, dynGENIE3 [85], does exist. However, we opted to utilize the original GENIE3 algorithm because we reasoned that the temporal spacing of developmental stages was likely too sparse for regulatory network reconstruction to incorporate time (note also that the co-expression algorithm we used, STEM, was explicitly designed for short time series such as ours). While our method therefore does not explicitly incorporate temporal dynamics, we purposefully biased our results to emphasize larval development over differences between replicates by only utilizing genes differentially expressed across larval development (or based on larval stage fed in the case of nurses).

We repeated the entire regulatory reconstruction reconstruction process 1000 times and averaged pairwise connection strengths across runs, as the algorithm is non-deterministic. To capture the total effect of each gene on the transcriptome dynamics within tissues, we averaged the regulatory effects each gene had on all other 999 genes expressed in the same tissue (“within-individual connectivity”). Similarly, to capture the effect each gene had on the transcriptome of social partners, we averaged regulatory effects each gene had on the 1000 genes expressed in social partners (“social connectivity”).

### Estimation of selective constraint, and evolutionary rate

Previously, we performed whole-genome resequencing on 22 diploid *M. pharaonis* workers as well as one diploid *M. chinense* worker to serve as an outgroup [48]. We estimated selective constraint using MKtest2.0 [86], assuming an equal value of alpha (an estimate of the proportion of nonsynonymous substitutions fixed by positive selection) across all genes. Selective constraint is the estimate of the proportion of nonsynonymous mutations that are strongly deleterious and thereby do not contribute to polymorphism or divergence [86]. Selective constraint is estimated using polymorphism data, so it represents the strength of purifying selection genes experience within the study population [54].

### Phylostratigraphic Analysis

Phylostrata are hierarchical taxonomic categories, reflecting the most inclusive taxonomic grouping for which an ortholog of the given gene can be found [87–90]. We focused on distinguishing between genes that were evolutionarily “ancient”, present in non-insect animals, versus genes present in only insects, hymenopterans, or ants [49]. We constructed a database containing 48 hymenopteran available genomes, 10 insect non-hymenopteran genomes, and 10 non-insect animal genomes (S2 Dataset). For outgroup genomes, we focused on well-annotated genomes which spanned as many insect orders and animal phyla as possible. Using this database, we estimated evolutionary age of genes based on the most evolutionarily distant identified BLASTp hit (E-value 10^−10^).

### Gene Set Enrichment Analysis

We performed gene set enrichment analysis based on social connectivity for each gene in each tissue separately using the R package topGO [91]. We identified enriched gene ontology terms using Kolmogorov-Smirnov tests (P < 0.05).

### General Analyses

We performed all statistical analyses and generated all plots using R version in R version 3.4.0 [92], aided by the packages “reshape2” [93], “plyr” [94], and “ggplot2” [95].

### Data Availability

All raw reads are available at DDBJ bioproject PRJDB3164. All source data for generating figures is included as S3 Dataset. All scripts and processed data (e.g. expression matrices, evolutionary measures) are available at https://github.com/warnerm/MonomoriumNurseLarva.

## Acknowledgments

We would like to thank the following: Mandy Tin for constructing RNA-sequencing libraries and performing RNA-sequencing, Luigi Pontieri for images of pharaoh ants, Chao Tong for compiling hymenopteran genomes for use in phylostratigraphy, and Rohini Singh for comments on the manuscript.

---

**S1 Fig.**
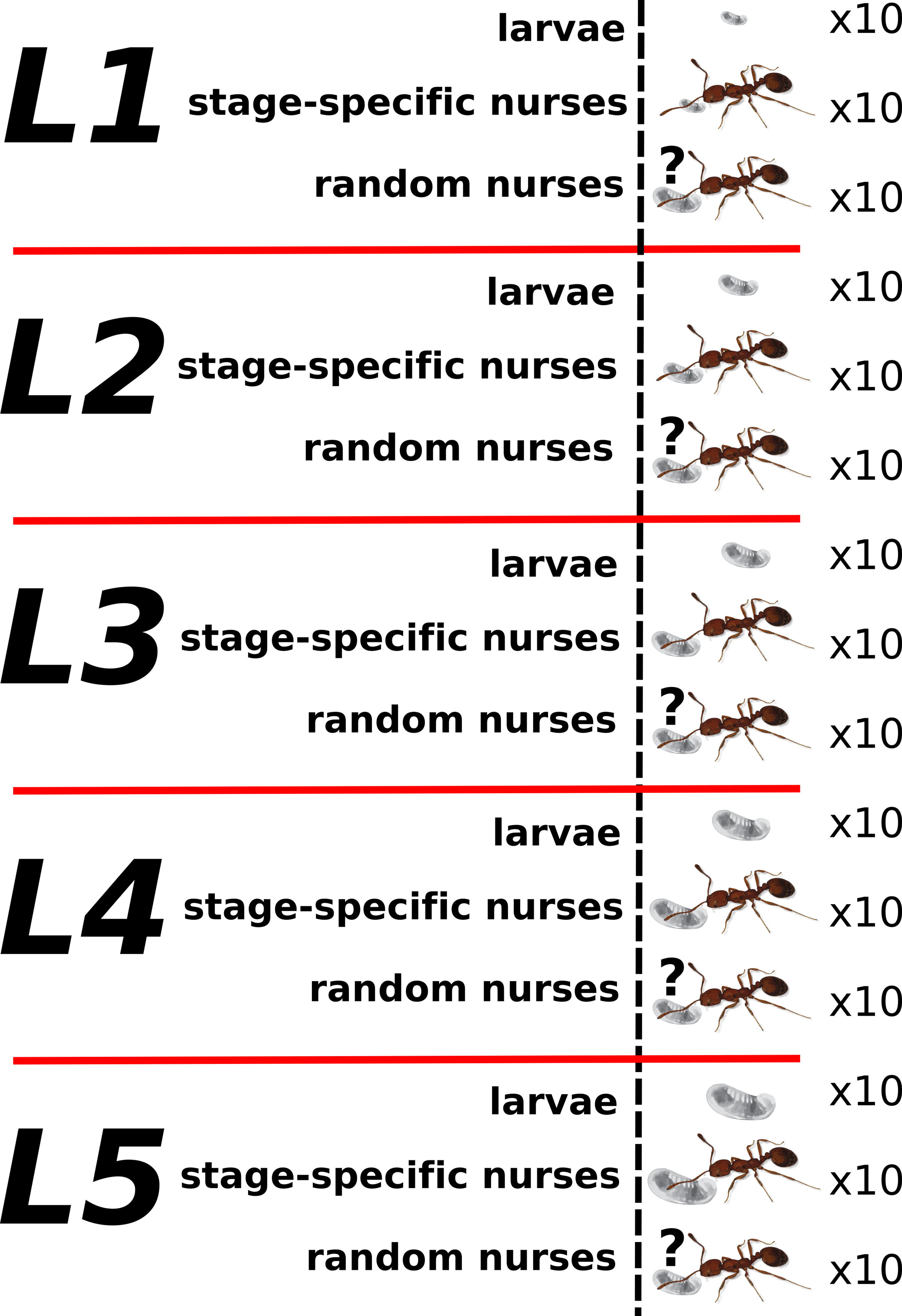
Diagram of sampling scheme. We collected ten worker-destined larvae, ten stage-specific nurses, and ten random nurses from each colony (six colonies per time point, where time points represent larval developmental stages L1, L2, etc). We collected stage-specific nurses when we observed them feeding larvae of the given developmental stage. We collected random nurses when we observed them feeding any stage of larvae.

**S2 Fig.**
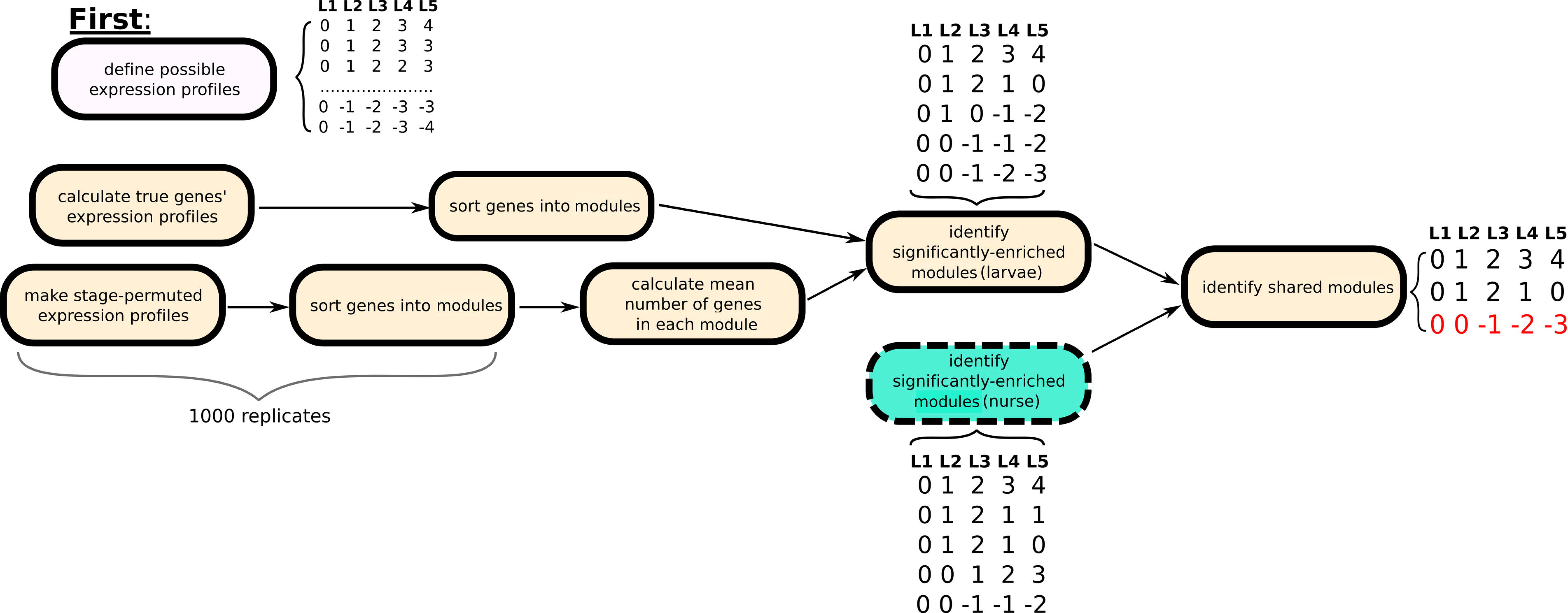
Identification of significantly-enriched modules shared between larvae and nurses. Inset tables depict pre-defined expression profiles of modules genes can be assigned to. First, we construct modules using all possible expression profiles (top left bubble). Expression profiles consist of five values, starting at zero, that indicate the log_2_ fold-change in expression from the initial value (at stage L1). At each subsequent stage, we either double, halve, or keep the expression level the same. This process is repeated to produce 81 (four stages after L1; 3*3*3*3 = 81) total modules. Next, for each tissue separately (here we depict workflow in larvae with yellow bubbles), we calculate individual gene expression profiles as the log_2_ fold-change in expression from the initial value at stage L1 and assign genes to the closest related module by Pearson correlation. Concurrently, we permute the developmental stage labels for each gene and assign the stage-permuted genes to modules (repeated 1000 times). From these stage-permuted results, we calculate the mean number of genes assigned to each module and treat this number as a null expectation (as each expression profile is not equally likely to occur by chance). We then identify significantly-enriched modules using a one-way binomial test (with the calculated mean as the null), with a Bonferroni-corrected false discovery rate of 0.05. This entire process is repeated in a nurse tissue and significantly-enriched modules are found (blue bubble). Finally, we compare significantly-enriched modules between larvae and nurses and retain identical and inverse modules as shared profiles. An example of an inversely related profile is shown in red, where larvae exhibit the enriched module [0, 0, −1, −2, −3] and nurses exhibit the inverse module, [0, 0, 1, 2, 3].

**S3 Fig.**
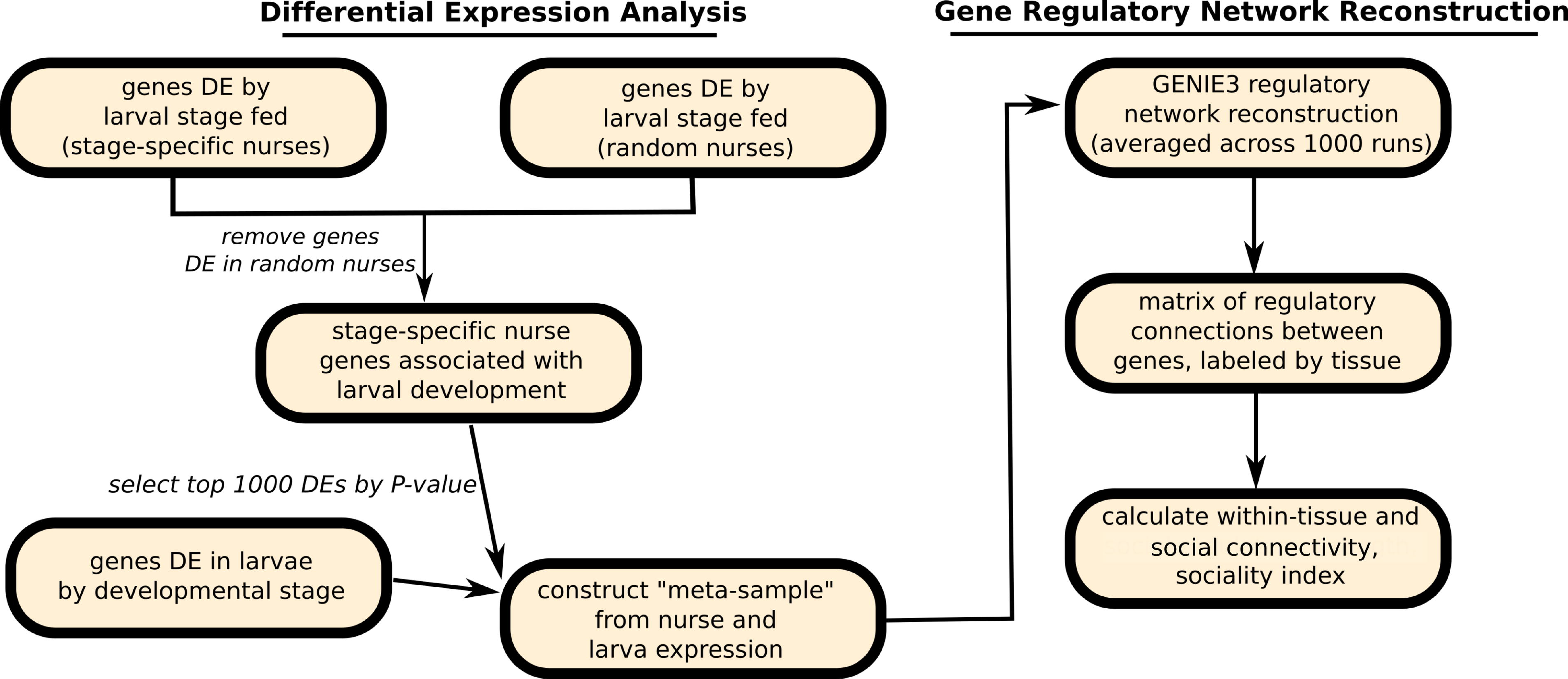
Workflow of preliminary differential expression analysis and gene regulatory network reconstruction. On the left, we identify putatively socially-acting genes through differential expression analysis. First, for nurse heads and abdomens separately, we perform differential expression analysis in stage-specific and random nurses to identify genes differentially expressed according to larval stage fed, using a nominal P-value of 0.05. We remove genes differentially expressed in random nurses, as these correspond to colony-specific environmental effects unrelated to social regulation of larval development. Next, we select the top 1000 differentially expressed genes by P-value in stage-specific nurses (after removing those DE in random nurses) as well as the top 1000 differentially expressed genes in larvae. From these genes, we create “meta-samples” by combining gene expression of larvae and stage-specific nurses collected from the same colony (separately for heads and abdomens), and labeling genes by the tissue they are expressed in. Using these meta-samples, we perform gene regulatory reconstruction (right) to identify genes expressed in nurses that regulate larval gene expression, and vise-versa. We repeat gene regulatory reconstruction 1000 times and average connection strength across runs, as the algorithm is non-deterministic. The output of gene regulatory reconstruction is a matrix of regulatory connections acting between genes. From this matrix, we calculate the average connectivity for each gene, separating within-tissue (larva-larva or nurse head-nurse head) from social (nurse-larva) connections. Genes with high connectivity are predicted to interact with many genes, i.e. are central to the network. Finally, we calculate each genes’ sociality index as the difference between social connectivity and within-tissue connectivity.

**S4 Fig.**
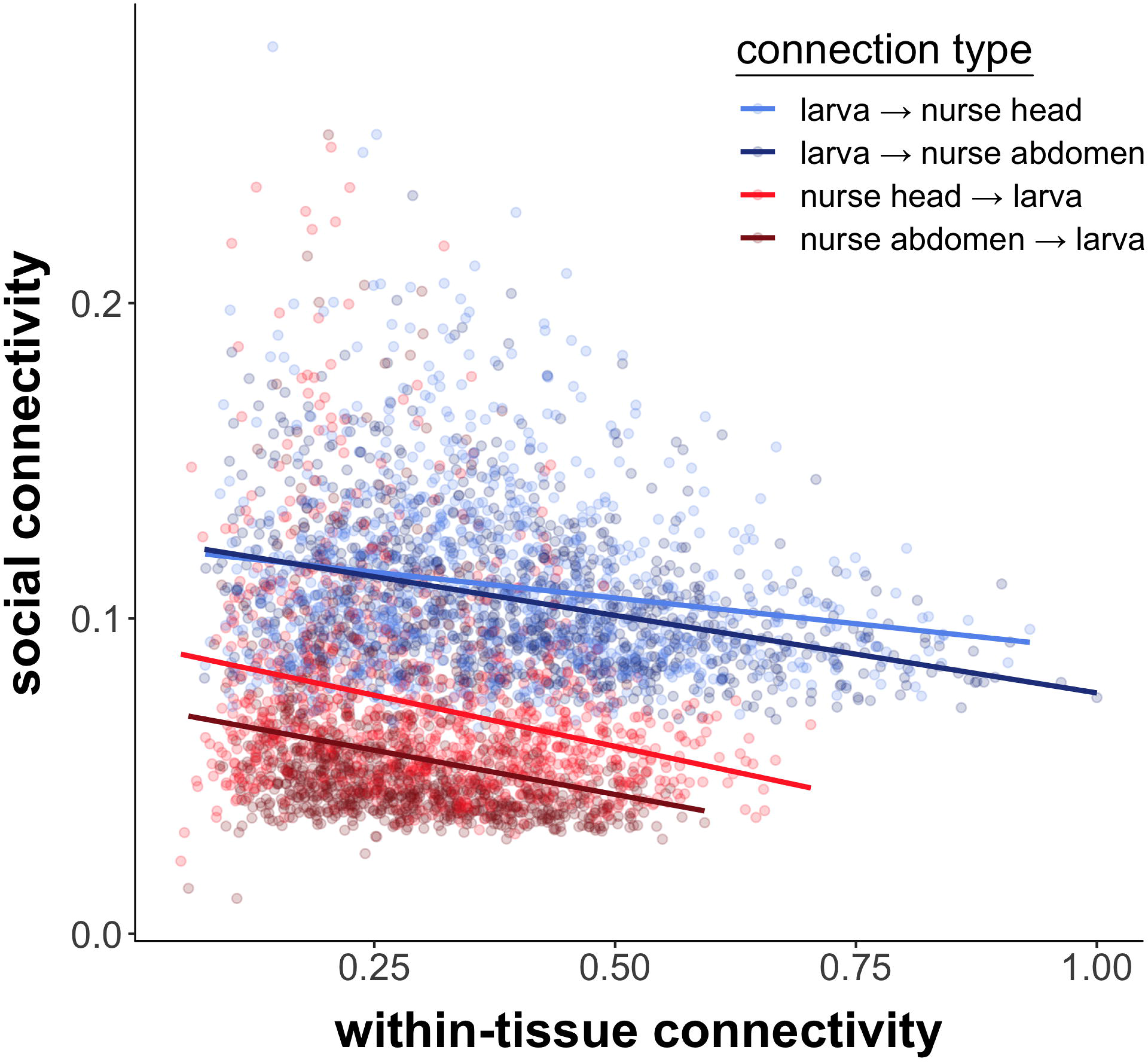
Genes highly connected in social regulatory networks are loosely connected in within-tissue regulatory networks. Connectivity is representative of the number and strength of regulatory connections each gene makes. Points indicate the average connectivity for a given gene, as measured within-tissue (x-axis; i.e. larva-larva or nurse-nurse) or socially (y-axis; i.e. larva-nurse). Points are colored by tissue the connectivity is measured in (e.g., dark blue indicates genes expressed in larvae, with connectivity measured in networks constructed with nurse abdomens). Spearman rho = −0.166, −0.374, −0.276, −0.342 for the four tissues as ordered in legend; P < 0.001 in all cases.

**S5 Fig.**
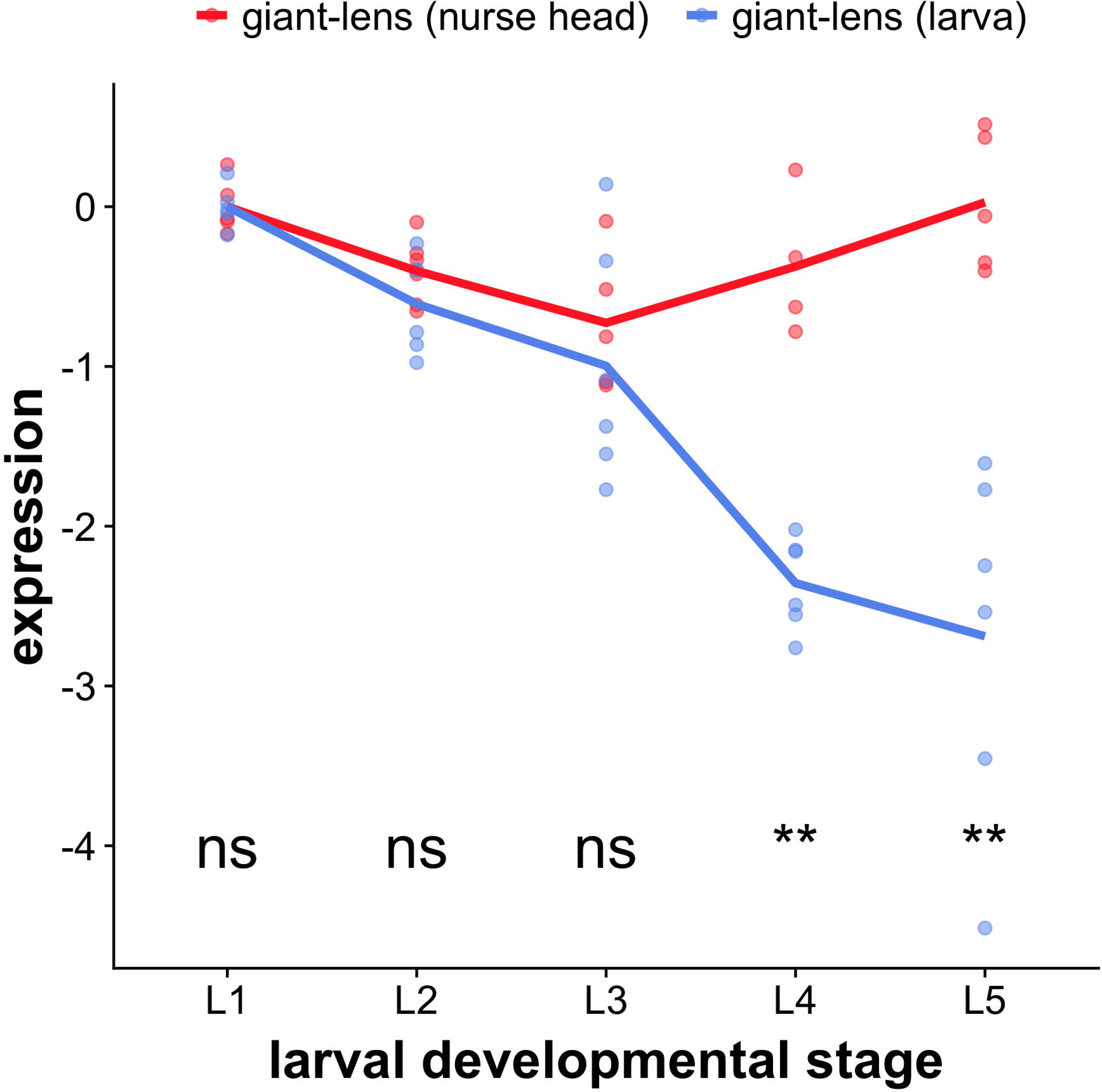
Expression of *giant-lens* in nurse heads and worker-destined larvae. Expression at stage *i* is equal to log_2_(expressioni/expression1), i.e. the ratio of expression at the given stage to expression at the initial (L1) stage. **: P < 0.01, ns: P > 0.05 (Wilcoxon test at each stage).

**S6 Fig.**
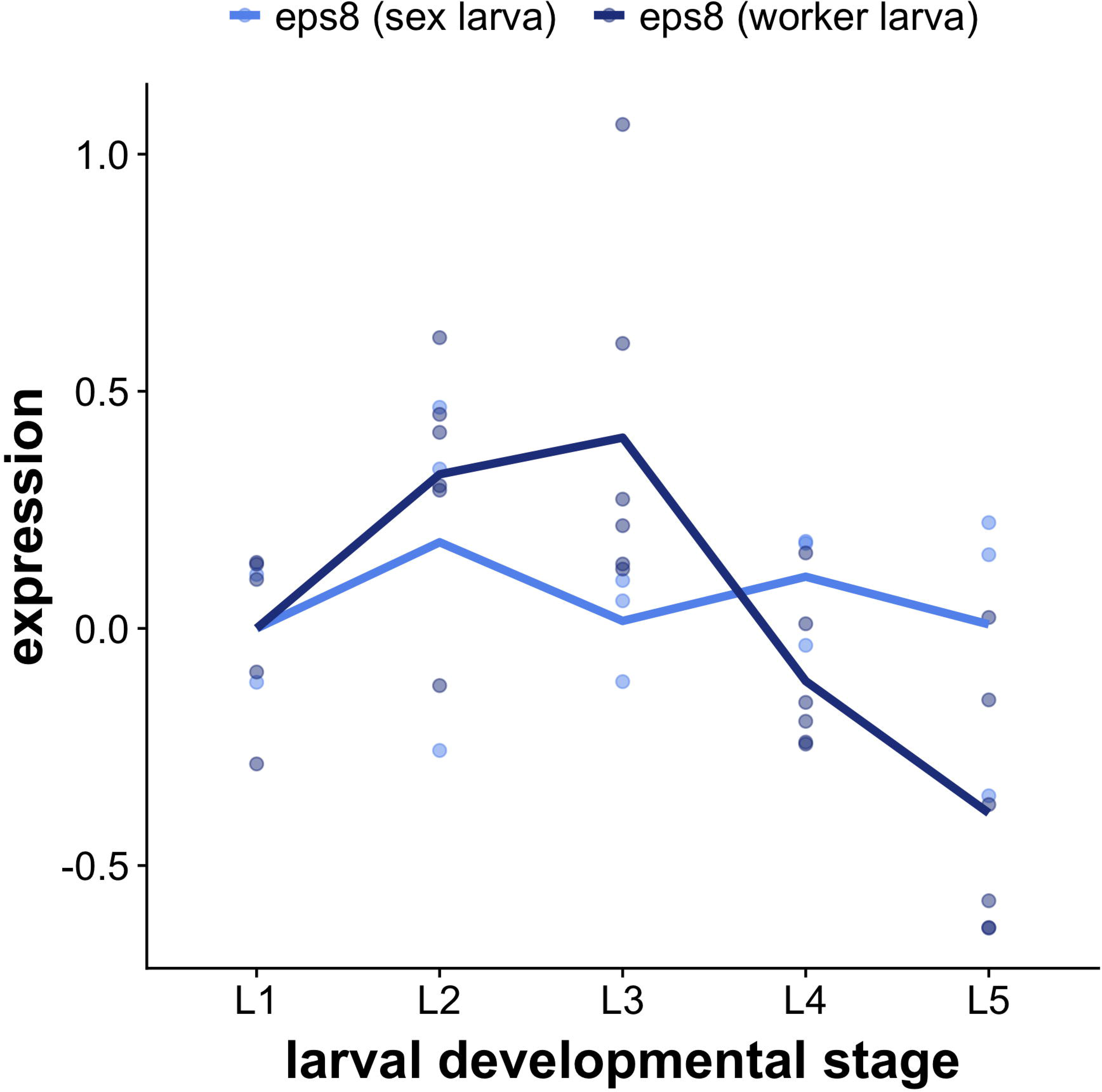
Expression of *eps8* (epidermal growth factor receptor substrate 8) in worker-destined and reproductive-destined larvae. Expression at stage *i* is equal to log_2_(expressioni/expression1), i.e. the ratio of expression at the given stage to expression at the initial (L1) stage. Expression of *eps8* changed differently over time in worker-destined versus reproductive-destined larvae (linear model with developmental stage treated as an ordinal variable; LRT; χ^2^ = 12.574, P = 0.014 for the interaction term stage*caste).

**Table S1.**
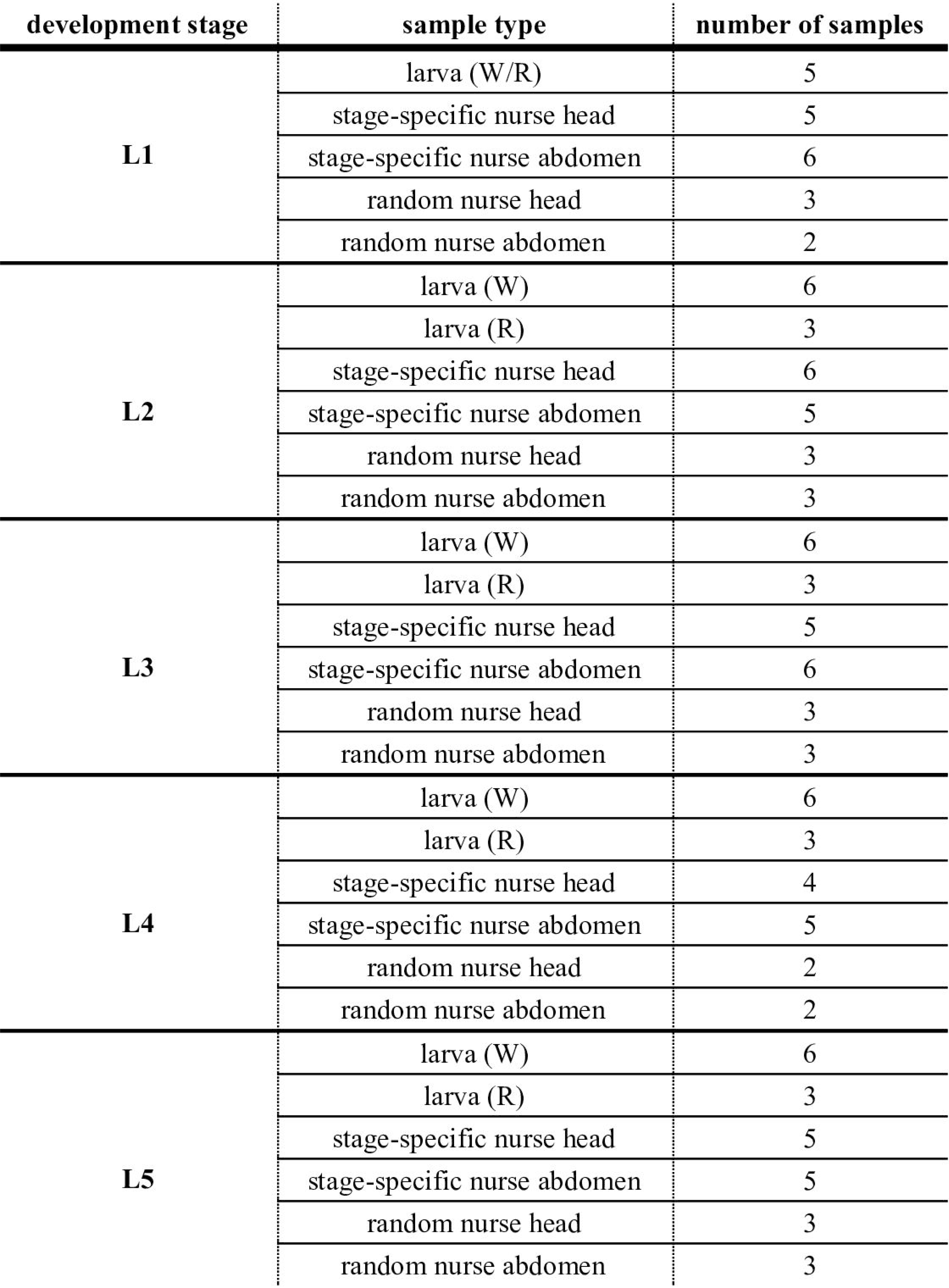
Description of samples included in study. Worker-destined larvae are indicated by larva (W), and reproductive-destined larvae are indicated by larva (R). Larval caste cannot be distinguished at the L1 stage, so L1 larvae are labeled larva (W/R). For network reconstruction, “meta” samples were used as input for network reconstruction, in which genes were labeled by sample type and grouped such that each gene contained a measurement of expression in worker-destined larvae, nurse heads, and nurse abdomens. After sample collection and RNA extraction, some samples exhibited clearly degraded RNA according to an Agilent Bioanalyzer assay. Removing these samples caused sampling to be uneven, so we used the minimum number of samples contained across tissues at a given stage for stage-specific nurse heads and abdomens, and randomly dropped excess samples. Overall, 25 “aggregate” samples were used as input for gene regulatory network reconstruction.

**Table S2.** Number of nurse significantly-enriched modules shared with larvae. Significantly-enriched modules are defined as modules with a statistically significant number of genes assigned, as determined by a permutation test (FDR < 0.05). Left column is the total number of significant modules for each tissue, while the second and third columns indicate number shared with larvae (out of 24 larval significantly-enriched modules). The last column indicates the total number of genes in these shared modules.

|  | number of<br>significantly-enriched modules | modules positively<br>shared with larvae | modules negatively<br>shared with larvae | number of genes<br>in shared modules |
| --- | --- | --- | --- | --- |
| *stage-specific nurse head* | 9 | 0 | 5 | 6838 |
| *random nurse head* | 9 | 0 | 2 | 209 |
| *stage-specific nurse abdomen* | 21 | 13 | 4 | 7943 |
| *random nurse abdomen* | 10 | 0 | 1 | 1400 |

**Table S3.**
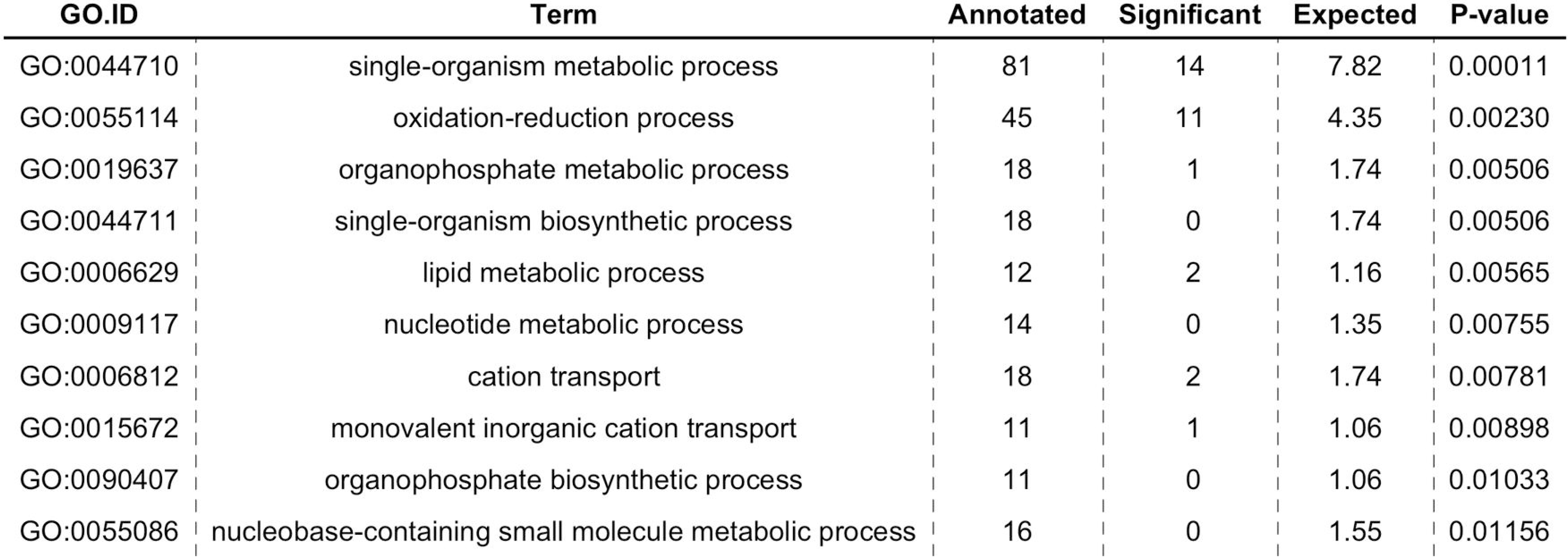
Nurse head social connectivity GO terms based on GSEA of social connectivity. P-value (unadjusted) is from Kolmogorov Smirnov (K-S) test. Enriched terms have higher than expected social connectivity in nurse heads.

**Table S4.**
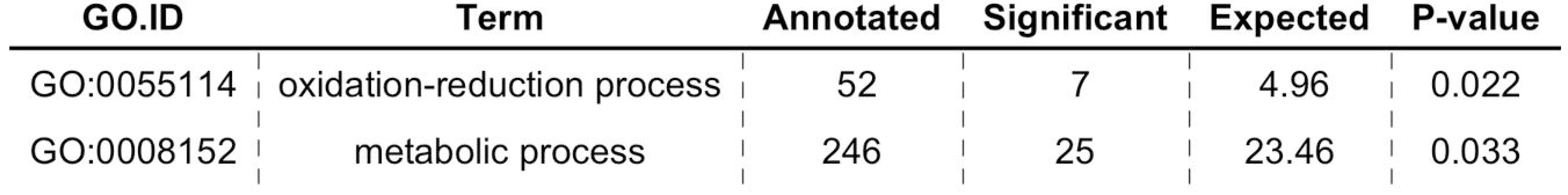
Nurse abdomen GO terms based on GSEA of social connectivity. P-value (unadjusted) is from Kolmogorov Smirnov (K-S) test. Enriched terms have higher than expected social connectivity in nurse abdomens.

**Table S5.**
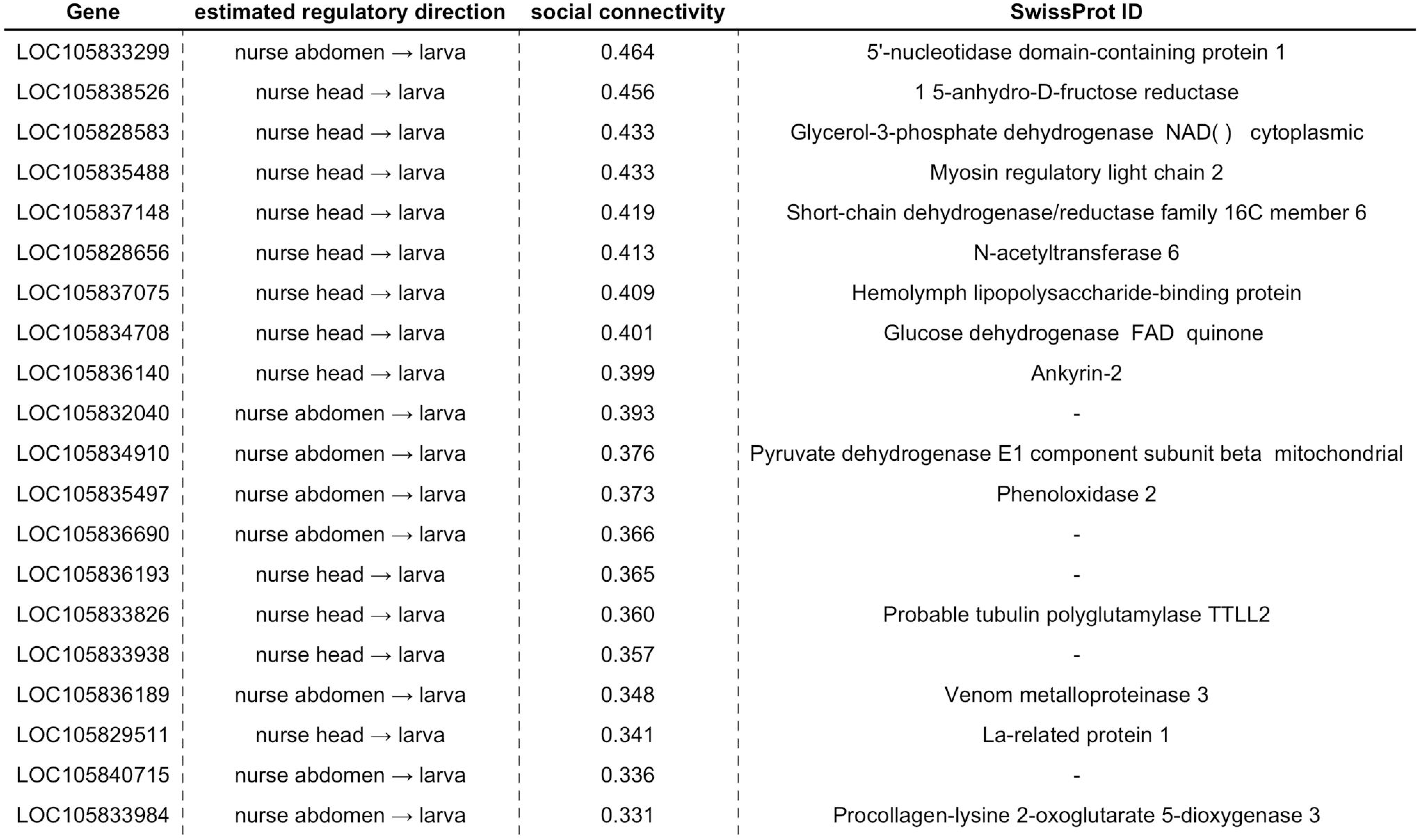
Top 20 genes by social connectivity in nurses. SwissProt ID is listed from automated annotation where a term was found.

**Table S6.**
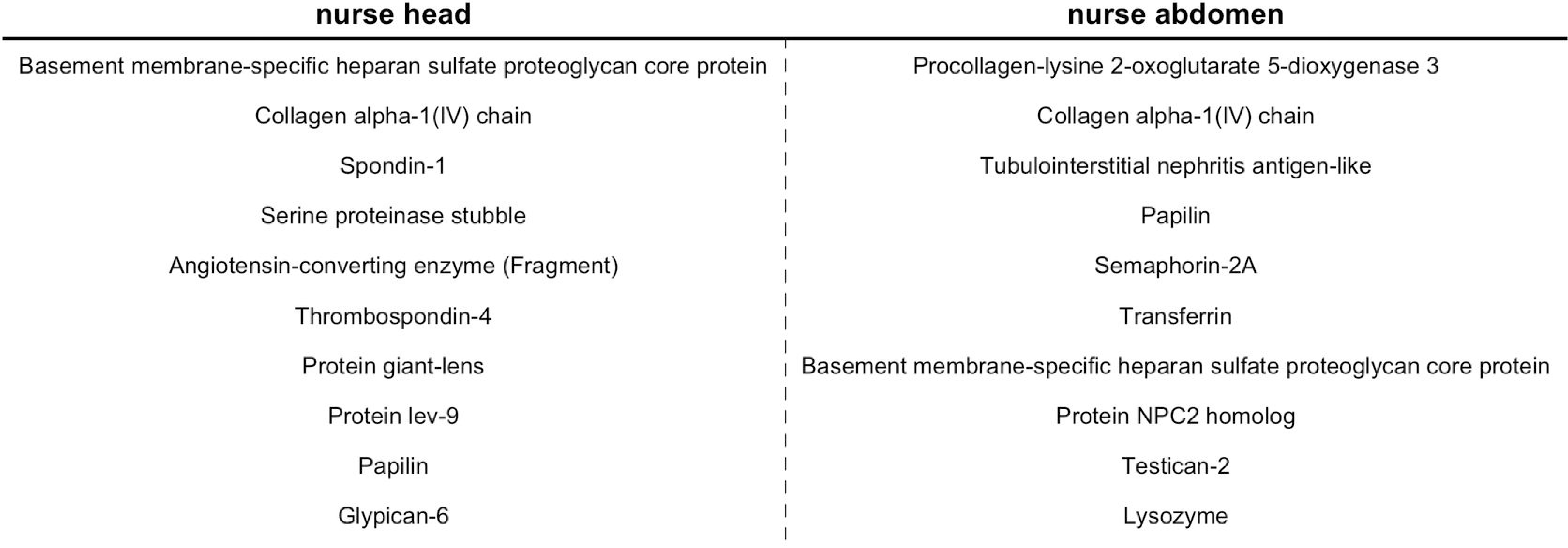
SwissProt annotations for the top genes coding for secreted proteins, sorted by social connectivity. Only genes with SwissProt annotations are included. All genes listed encode for secreted proteins in *D. melanogaster*.

| nurse head | nurse abdomen |
| --- | --- |
| Basement membrane-specific heparan sulfate proteoglycan core protein | Procollagen-lysine 2-oxoglutarate 5-dioxygenase 3 |
| Collagen alpha-1(IV) chain | Collagen alpha-1(IV) chain |
| Spondin-1 | Tubulointerstitial nephritis antigen-like |
| Serine proteinase stubble | Papilin |
| Angiotensin-converting enzyme (Fragment) | Semaphorin-2A |
| Thrombospondin-4 | Transferrin |
| Protein giant-lens | Basement membrane-specific heparan sulfate proteoglycan core protein |
| Protein lev-9 | Protein NPC2 homolog |
| Papilin | Testican-2 |
| Glypican-6 | Lysozyme |

**S1 Dataset. Complete list of all differentially expressed genes.**

Each gene can be differentially expressed in three tissues: worker larva, nurse head, or nurse abdomen. P-values are listed for each tissue. The top 1000 differentially expressed genes (by P-value) were used for regulatory network reconstruction. Social connectivity is the sum of all regulatory interactions in the direction specified by “estimated regulatory direction”.

**S2 Dataset. List of species used for phylostratigraphy.**

Each species listed, with their NCBI taxonomic ID, was used in the construction of the phylostratigraphic database to estimate evolutionary ages of genes.

**S3 Dataset. Data files used to construct all figures.**

All data are organized in text files, with the relevant figure listed in the title.

